# TCID50 Measurements of anti-viral efficacy on metal printed masks and plastic surfaces

**DOI:** 10.1101/2022.10.13.512105

**Authors:** Colette S.M. Bilynsky, Kishana Taylor, Megha Anand, Elizabeth Wayne

## Abstract

The SARS-CoV-2 pandemic has created a need for effective personal protective equipment (PPE) to prevent viral spread. PPE like face masks contain the spread of virus-filled droplets and thus reduce infection rates, has been a critical tool in stopping the spread of SARS-CoV-2. PET plastic barriers have also been used in public settings to reduce face to face viral transmission. However, in some cases, they have provided additional contact with the virus due to contamination. In order study, we evaluated the effectiveness of face masks and PET plastics coated in different metals in reducing viral load. We compared PPE printed with silver, copper, or zinc for their ability to inactivate live human coronavirus HCoV 229E. Our results show that silver and copper have significant anti-viral efficacy when printed on nonwoven fabric compared to the controls. The metal-printed PET showed around 70% anti-viral efficacy with any formulations, with copper performing the best. This work builds more data to support the development of metal printed materials for enhanced protection against coronaviruses.

## Introduction

According to the World Health Organization report compiled at the time of this writing, the Severe Acute Respiratory Syndrome coronavirus 2 (SARS-CoV-2) has resulted in 528,816,317 cases worldwide and nearly 6.3 million deaths (1). More than twelve variants have been identified, each with varying levels of infectivity and resultant COVID19 disease severity. SARS-CoV-2 transmission is respiratory, found in droplets, and fine particle aerosols, shed by infected persons. Implementation of non-pharmaceutical interventions, such as-social distancing, eye protection, and mask-wearing, have been implemented to reduce contact with virus particles.

Protective face masks such as cloth masks, surgical masks, and N95 medical respirators have been pivotal in reducing SAR-CoV-2 population spread (2). In addition, face mask usage has resulted in a reduction of other respiratory illnesses like Influenza (3). Face mask type, user fit, and adherence to proper use vary significantly and can impact efficacy. Surgical masks are also recommended for protection against SARS-CoV-2 infection and tend to be made from a thin plastic material, polypropylene, and are disposable. Cloth masks can be comprised of many fabric types (i.e., cotton, silk, tea towel, linen) and are a sustainable option because they can be washed and reused (5). Cotton masks and surgical masks were also found to decrease viral uptake, albeit to lesser extents, exhibiting a 20-40% reduction for cotton masks and a 50% reduction for surgical masks (4). Cloth masks improved in efficiency with increased threading density (6) or with increased layers (7). This is hypothesized due to mechanical and electrostatic filtration effects (7). N95 masks have been shown to decrease viral uptake by 80% if generally worn and by 90% if fitted for a proper seal compared to non-masked scenarios (4). It must be noted that some systematic reviews show non-definitive results in the efficacy between N95 and surgical masks (8). However, this study analyzed papers from 1990-2014 (8) or from early 2020 (9) and may not account for transmission differences from more infectious SARS-CoV-2 variants such as Omicron. However, none of these mask-were able to ultimately impede exposure to viral particles, hindering their effectiveness (4)

Polyethylene Terephthalate (PET) is used to make face shields and barriers often used in medical and commercial settings (10). SARS-CoV-2 is known to persist on various surfaces for several days (11,12), remaining active on surgical masks for as long as six days (13). This raises concerns about indirect transmission through contact with contagious Personal Protective Equipment (PPE). Previous studies have indicated that 7% of confirmed COVID-19 cases in the healthcare personnel population have been attributed to contact with contaminated surfaces and PPE (13). As such, there is likely a similar portion of general population cases caused by this route of exposure. This risk of indirect transmission coupled with viral penetration of pre-existing masks, therefore, necessitates the development of masks and other PPE with anti-viral properties to either kill or inactivate incident viruses.

Metals with antibacterial (14) and anti-viral properties (15) have been used as a synergistic enhancement to PPE. Metals can have anti-viral activity via disrupting virus membranes and interaction with virus replication factors (16). Recent studies have also shown that certain metals, mainly copper, silver, and zinc, were effective in inactivating and reducing surface viral load (17). Copper has been mechanistically shown to exhibit anti-viral properties. The mechanism for this involves stimulating the generation of Reactive Oxygen Species (ROS), damaging viral DNA and proteins (13,18). Silver nanoparticles exhibit anti-viral properties by preferentially binding to viral surface proteins, disrupting its exterior interactions, and modifying protein structure to destabilize and inactivate the virus (19). Zinc has been shown to inhibit the viral RNA Polymerase activity, preventing replication (20,21).

Metal printed materials have also shown efficacy against other viruses such as enveloped viruses like severe acute respiratory syndrome (SARS) coronavirus, Human Immunodeficiency Virus (HIV), Influenza (22), and Dengue (23). These materials have also been used to destroy non-enveloped viruses like norovirus (24) and infectious bursal disease virus (25), which are more resistant to drying on inanimate surfaces (26,27) and anti-viral therapy (28,29). Tremiliosi et al. (2020) created silver nanoparticle polycotton fabric (30). Brokow et al. (2010) designed anti-influenza face masks infused with copper oxide (31). Hamouda et al. (2021) even created wipes with silver nanoparticles that were effective against coronavirus (32).

The wide variety of techniques and metals used to create anti-viral and anti-microbial materials make it imperative to have studies that compare multiple formats. In this study, we investigate the anti-viral efficacy of metal incorporation into commonly used PPE materials, non-woven face masks and PET plastics. We are compare the anti-viral efficacy of silver, copper, and zinc on both fabric and PET using the Reed and Munch Method to determine the TCID50 dose.

## Materials and Methods

### Viruses and Virus Titers

Human coronavirus 229E (HCoV 229E) was purchased from American Type Culture Collection (ATCC, Manassas, VA). HCoV 229E virus was inoculated onto MRC5 cells (ATCC) and Huh7 (Sexisui Xenotech, Kansas City, KS) cells to generate virus stocks. Approximately 100 to 1000 μL of the virus was inoculated onto monolayered cells and watched for cytopathic effect (CPE) over a period of 48 hours for MRC5 cells and 30 hours for Huh7 cells. The supernatant was collected, spun down to remove any floating cells, and endpoint titration was performed to ascertain the median tissue culture infectious dose (TCID50). Briefly, 100 μL of HcoV 229E supernatant was diluted in 900μL of media for eight serial dilutions. Each dilution was serially plated in 96-well plate format. Wells were monitored for CPE for five days, and the Reed and Munch method was used to calculate the viral titers as 10^x^ TCID50 /mL. All experiments with the live virus were conducted in EH&S approved BSL 2 facilities using a dedicated cell incubator for all viral experiments.

### Cell Culture

MRC5, lung fibroblast cells (ATCC, Manassas, VA), were cultured in Eagle’s Minimum Essential Medium (EMEM) supplemented with 10% FBS and 1% penicillin: streptomycin. Huh7, hepatocarcinoma cells (Sexisui Xenotech, Kansas City, KS) were cultured in DMEM supplemented with 10% FBS and 1% penicillin: streptomycin. All cells were incubated at 37 C and 5% CO2. MRC5 cells were used within ten passages throughout the experiments.

### Cytotoxicity

Cells were seeded in 96-well plates and allowed to adhere overnight. Mask materials were vortexed with media and then added to the cells to incubate for two hours. Twenty-four hours following exposure to the material, cell viability by measured by Dojindo Cell Counting Kit-8 (CCK-8) using the manufacturer’s protocol. Percentage viability was found by comparing treated cells to untreated controls.

### Face Masks and PET materials

Fabric and PET coated with copper, silver and zinc materials were obtained from Liquid-X (Pittsburgh, PA). Uncoated control fabric and PET materials were obtained from the same company. The surgical and cotton masks measured in this study were sourced from commercial vendors.

### Virus Exposure

Stock HCoV 229E virus was inoculated onto each sample (fabric and PET) and allowed to incubate for 1-2 hours. Media was added to the samples, and endpoint titrations to the −5 were performed to assess the viral load remaining on each sample. After five days of observation, wells exhibiting cytopathic effects were marked, and the titer was calculated using a Reed and Munch table. Samples were treated in duplicate. A back titer of the viral stock was performed with each experiment. The log reduction was calculated using the R-log reduction method (33–35). C_a_ is the TCID50 value of the infected group, and Cb is the TCID50 value of the uninfected control group.

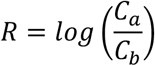

### Statistical Analysis

Figures were plotted using GraphPad Prism (San Diego, California), and one-way ANOVA followed by post-hoc Tukey tests were performed to assess significance, except in Figure 6, where an unpaired t-test was performed. Significance is denoted with asterisks correlating to P values as follows: p<.0001(****), p<.001 (***), p<.01 (**), p<.05(*). Microsoft Excel (Redmond, Washington) was used to find the standard deviation and mean.

## Results

### Measurement of cytotoxicity of anti-viral PPE

To demonstrate the efficacy of anti-viral coating against SAR-CoV-2 viral spread, we inoculated PPE materials against human coronaviruses **(Figure 1)**. We used HCoV 229E an alpha human coronavirus as a surrogate for SARS-CoV-2 because it is BSL2 compliant and is one of the four common coronaviruses, contributing to 15%-30% of common cold cases (36). Viral titers are traditionally formed using a plaque assay. Since HcoV 229E viral infection does not result in plaque formation, we measured viral titers using the Reed and Munch method rather than a traditional plaque assay. The Reed and Munch method calculated by measuring the degree of cytopathic effect (CPE) on infected cells.

**Figure 1.**
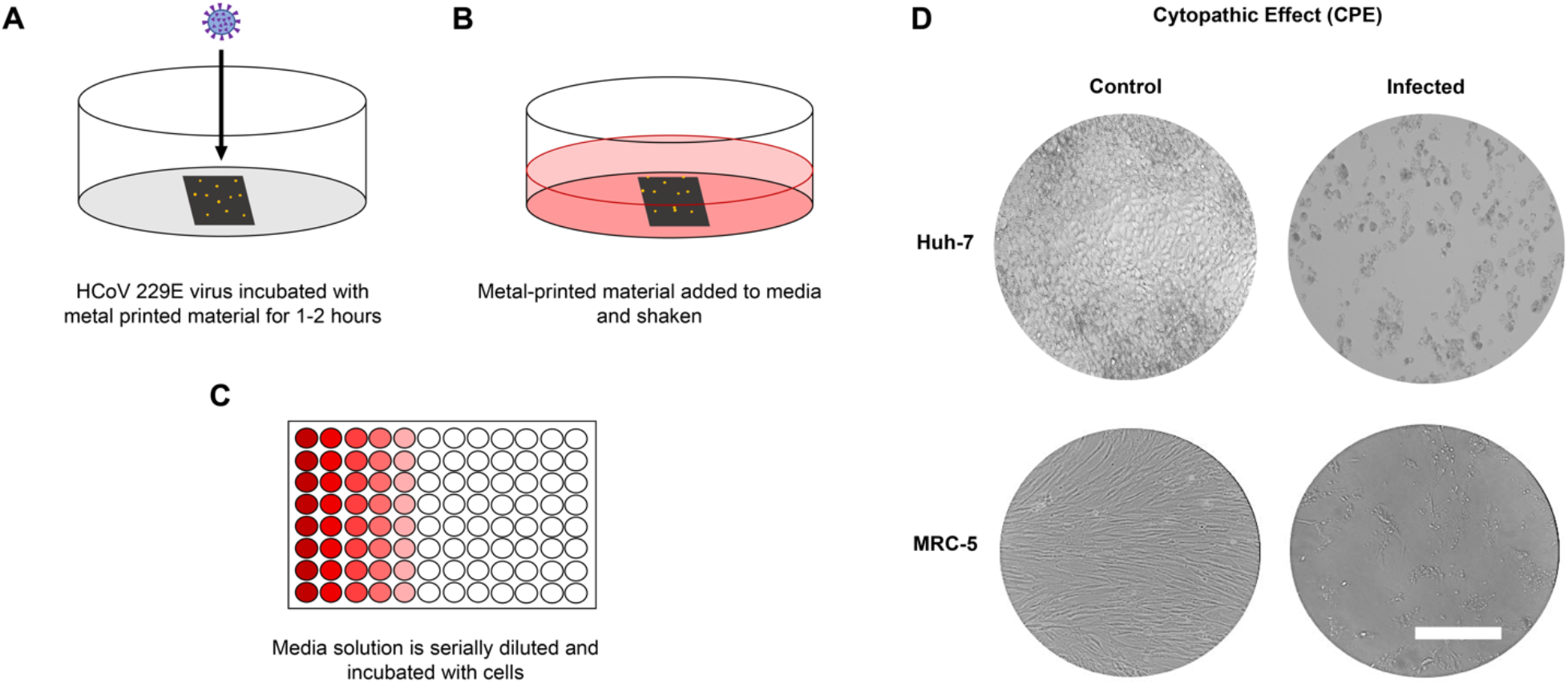
TCID50 Endpoint assay to assess anti-viral efficacy on personal protective equipment. **A)** HCoV 229E virus is incubated with metal printed materials for either 1 or 2 hours. B) Metal printed material is then added to complete cell culture media and shaken to remove the remaining free virus from the material. C) Media is serially diluted into wells pre-incubated with either Huh-7 or MRC-5 cells following the TCID50 Reed and Munch assay. D) Bright-field images of Huh-7 liver cells (top) and MRC5 human lung fibroblast cells (bottom) immediately before and five days after HCoV 229E infection. Scale bar = 200um.

### Silver and Copper-printed fabric show a reduction in viral titer

Before using live virus, we first tested the effect of the metal printed materials on cell viability using a Cell Counting Kit (CCK-8) assay **(Figure 2A and 2C).** The zinc and silver-printed nonwoven fabric did not decrease the viability of the MRC5 cells nor the Huh7 cells. However, the copper printed mask drastically reduced MRC-5 viability. In contrast, the Huh-7 cells showed little toxicity when exposed to copper-printed masks. However, exposure to copper-printed masks resulted in a 64.1% (± 3.9%) viability in the Huh7 cells and an 11.1% (± 5.4%) viability in MRC5 cells. In addition, the combination, silver-copper-printed masks showed a significant cytotoxic effect compared to the controls in both cell lines. Huh7 cells exposed to the silver-copper-printed fabric had a 20% (± 5.5%) viability, while MRC5 cells had a 2.3% (±5.9%) viability. Thus, we investigated the effects of copper-printed masks with Huh-7 cells. For the other masked groups, we studied anti-viral activity using both cell lines.

**Figure 2:**
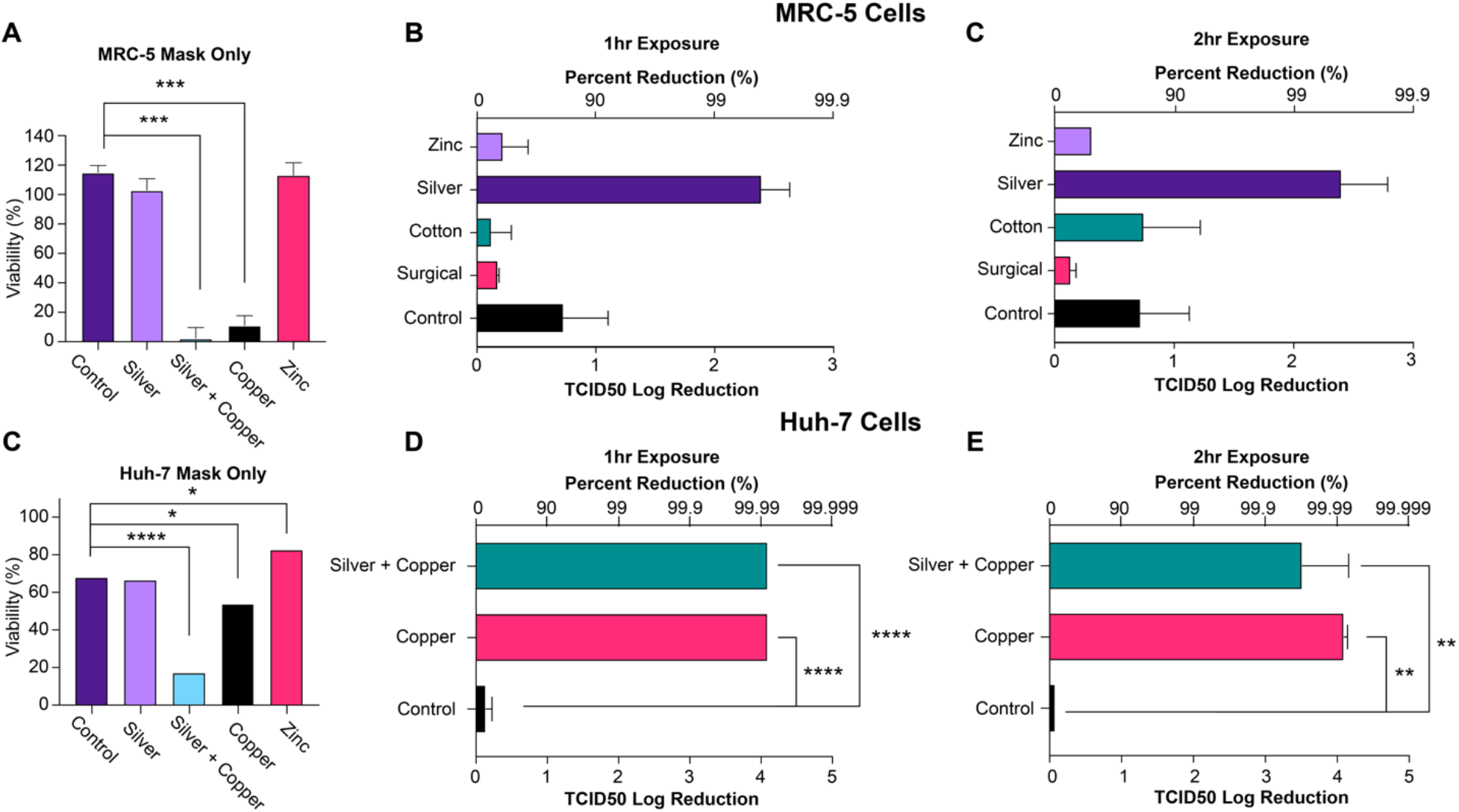
Silver shows the highest reduction in viral load among metal printed mask candidates. CCK8 viability assay of metal printed masks on **A)** MRC-5 cells and **B)** Huh-7 cells. All cells were normalized to the cells not exposed to masks. zinc nonwoven, silver nonwoven, control nonwoven, cotton cloth masks, and medical/surgical masks were exposed to HcoV 229E live virus before incubation with cells. TCID50 was calculated after **C)** 1 hour or **D)** 2 hours for MRC5. One-way ANOVA tests were performed. *p<0.05, **p<0.01, ***p<0.001, **** = p<0.0001.

Several metals were tested for their anti-viral effectiveness **(Table 1).** We found that silver-printed fabric demonstrated a significant reduction in viral titer in comparison to the controls and other metal-printed materials. At 1 hour, there is a 4.09 (± 0.035) log reduction after exposure to either the copper-printed or silver-copper printed fabric compared to the 0.139 (± 0.06) reduction seen after exposure to the control fabric. Notably, there was no significant decrease in viral titer between 1-hour and 2-hour exposure times amongst any of the metal-printed groups. Within the non-printed masks examined, the exposure time reduced viral titer. However, this might be due to the instability of the viral under the experimental conditions rather than any anti-viral activity (***Figure 2***).

**Table 1:** Chart of materials and their descriptive properties.

| Control Nonwoven | Uncoated nonwoven fabric control for masks |
| --- | --- |
| Silver Nonwoven | Silver-printed nonwoven fabric |
| Silver-Copper Nonwoven | Silver-and-Copper-printed nonwoven fabric |
| Copper Nonwoven | Copper-printed nonwoven fabric |
| Zinc Nonwoven | Zinc-printed nonwoven fabric |
| Cotton | Cotton from a reusable mask |
| Surgical | Surgical mask material |
| Control PET | Uncoated PET |
| Copper PET | Copper-printed PET |
| Zinc PET | Zinc-printed PET |

### Copper-printed PET material causes a significant reduction in viral load compared to control and zinc-printed PET

Three different types of PET material were tested for efficacy in reducing viral titer (*Table 1*). Copper and zinc printed PET were tested on MRC5 and Huh7 cells with varying results, as seen in **(Figure 3)**. Copper did not induce cytotoxicity as in the case of the printed masks; however, the zinc did show toxicity. At 1 hour, the virus exposed to the copper-printed PET had a 0.426 (± 0.045) log reduction when tested on the MRC5 cells (compared to a 0.079 log reduction on the control PET) and a 0.274 (± 0.035) log reduction on the Huh7 cells which is slightly less than the reduction seen by the control PET (0.309). This same trend was seen at 2 hours. The copper-printed PET also performed significantly better than the zinc-printed PET at 1 hour, which only had a log reduction of 0.129 (± 0.05) in viral titer.

**Figure 3:**
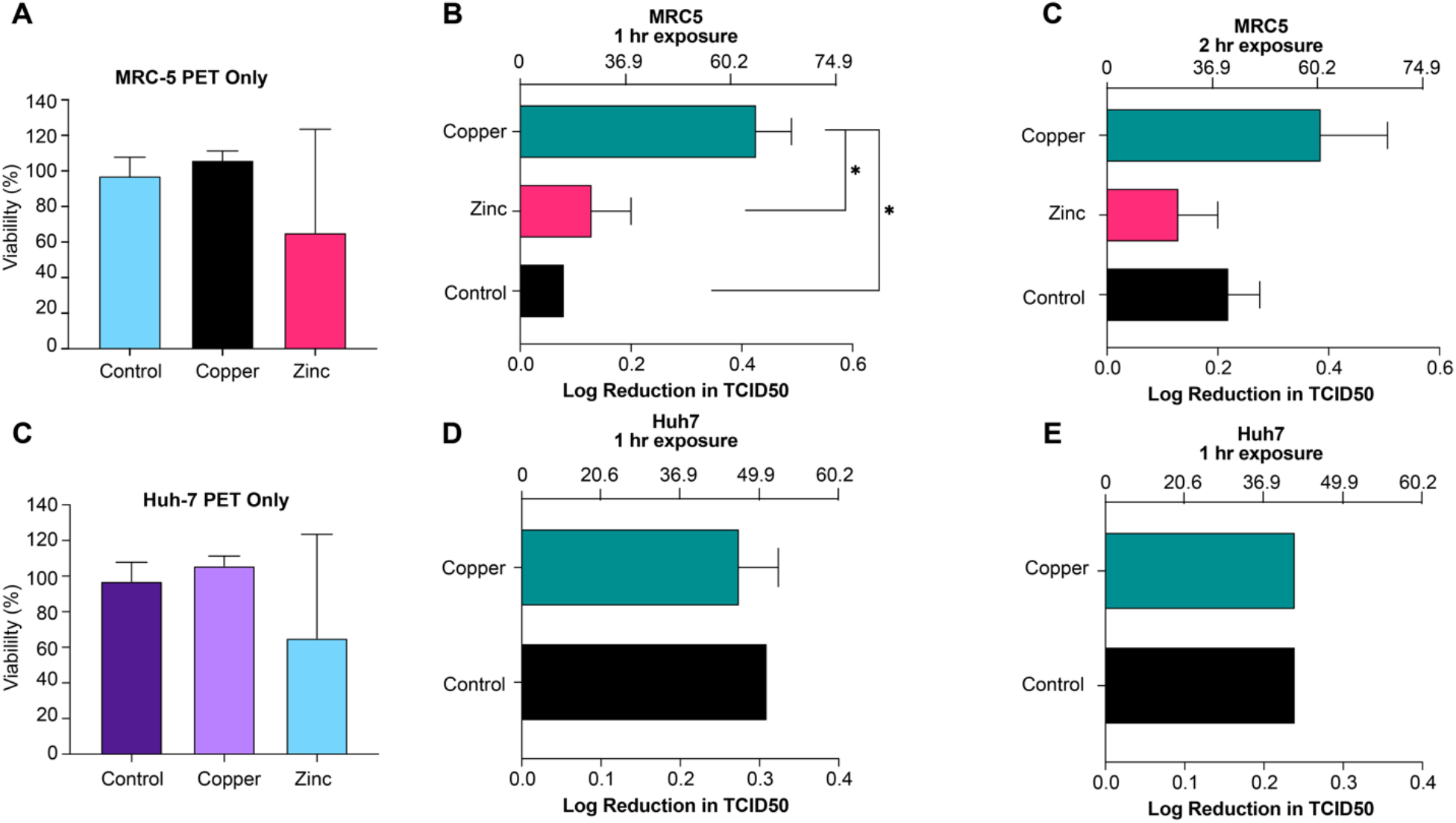
Copper shows the highest reduction in viral load among the PET plastic candidates. CCK8 viability assay of metal printed PET plastics on **A)** MRC-5 cells and **C)** Huh-7 cells. All cells were normalized to the cells not exposed to masks. Zinc nonwoven, silver nonwoven, non-printed (control) PET was exposed to HcoV 229E live virus before incubation with cells. TCID50 was calculated after **C)** 1 hour or **D)** 2 hours for MRC5. One-way ANOVA tests were performed. * = p<0.05.

## Discussion

In this study, we tested both metal-printed fabric for the use of face masks as well as metal-printed PET for barriers and other high-touch surfaces. We aimed to calculate the effectiveness of different metals in reducing viral load when they encounter HCoV 229E as a surrogate for SARS-CoV-2. Because copper was shown to be cytotoxic to MRC-5 cells, we used Huh7, which has been shown to be permissive to HCoV 229E infection, to test the efficacy of copper in reducing the viral load (39). However, this did result in some variable results between cell lines, suggesting differences in cell line sensitivity.

Silver-printed masks were found to be the most effective at 1 and 2 hours when tested on the MRC5 cell lines. This aligns with plentiful evidence that silver has potent anti-viral abilities, especially for SARS CoV-2 (17,19,30,32,38). Similarly, we found that the copper and silver-copper printed fabric and PET caused significant decreases in viral titer. Copper is another metal that has demonstrated anti-viral and antimicrobial abilities (13,17,18,31). The zinc-printed fabric and PET both did not display a significant antiviral effect. Zinc has been shown to have some anti-viral abilities (20,21,37), that were not corroborated in this study, perhaps due to differences in formulation or concentration of zinc utilized. We expected to see an increased log reduction after the second hour based on results published in previous literature (11). However, we did not see this increased reduction with the second hour of exposure.

Our results regarding the anti-viral efficacy of copper have limitations mostly due to the cytotoxic effect of copper. While we did not observe a significant cytotoxic effect in the metal-printed PET materials, the metal-printed fabrics did show some cytotoxic effect to both the Huh7 and MRC5 cell lines. Both the copper and silver-copper printed fabric were found to be significantly more cytotoxic than the control fabric and the silver-printed fabric. Previous literature has found copper to be more cytotoxic than silver, even though it does carry advantages in expense and abundance (40). Interestingly, this trend of copper cytotoxicity was not seen in the PET material, which may be due to differences in the porosity of the material or printing method. Metal-infused fabrics have been shown to leech metals after exposure to water or saliva (41), which could account for copper-printed fabrics decreases in cell viability. Our results did not demonstrate any cytotoxicity from the zinc-printed materials, somewhat at odds with previous studies that saw dose-dependent cytotoxicity of zinc oxide in MRC5 cells (42).

## Conclusion

PPE is an effective way to reduce exposure to SARS-CoV-2 as well as other respiratory viruses. Masks are very effective in reducing. However, even N95 masks, even when fitted properly, have resulted in some transmission. (4). High-touch areas like healthcare centers, public spaces, and homes were also potential sources of transmission (37). While frequent disinfection can eliminate surface contamination, making them anti-viral by design would help limit transmission. Coating with anti-viral materials like metal can add additional protection by killing viral particles on PPE surfaces (37). This study determined that copper and silver printed masks and PET surfaces can reduce the viral titer after only 1 hour of exposure. Further studies to evaluate its effectiveness in daily life usage should be explored.

## List of abbreviations

COVID-19: Coronavirus Disease 2019
SARS-CoV-2: Severe Acute Respiratory Syndrome coronavirus 2
PPE: Personal Protective Equipment
ROS: reactive oxygen species
DNA: Deoxyribonucleic acid
RNA: Ribonucleic acid
SARS: severe acute respiratory syndrome
HIV: Human Immunodeficiency Virus
PET: Polyethylene Terephthalate
TCID50: median tissue culture infectious dose
HCoV-229E: Human coronavirus 229E
MRC5: Medical Research Council Cell Strain-5
Hr: Hour
ACE2: Angiotensin converting enzyme 2
ATCC: American Type Culture Collection
FBS: fetal bovine serum
EMEM: Eagle’s Minimum Essential Medium
DMEM: Dulbecco’s Modified Eagle Medium
CCK-8: Cell Counting Kit-8

## Authors’ contributions

CSMB performed experimental design, conducted experiments, performed analysis, and wrote the manuscript. KT performed experimental design and experiment analysis. MA conducted experiments and performed analysis. ECW performed experimental design and analysis and wrote the manuscript.

## Acknowledgments

We would like to thank LiquidX Metals, Inc for providing the masks and PET materials used in this study.

## References

1. WHO Coronavirus (COVID-19) Dashboard [Internet]. [cited 2022 May 26]. Available from: https://covid19.who.int

2. Liang M, Gao L, Cheng C, Zhou Q, Uy JP, Heiner K, et al. Efficacy of face mask in preventing respiratory virus transmission: A systematic review and meta-analysis. Travel Medicine and Infectious Disease. 2020 Jul 1;36:101751.

3. Sande M van der, Teunis P, Sabel R. Professional and Home-Made Face Masks Reduce Exposure to Respiratory Infections among the General Population. PLOS ONE. 2008 Jul 9;3(7):e2618.

4. Ueki H, Furusawa Y, Iwatsuki-Horimoto K, Imai M, Kabata H, Nishimura H, et al. Effectiveness of Face Masks in Preventing Airborne Transmission of SARS-CoV-2. mSphere. 2020 Oct 28;5(5):e00637–20.

5. Lima MM de S, Cavalcante FML, Macêdo TS, Galindo-Neto NM, Caetano JÁ, Barros LM. Cloth face masks to prevent Covid-19 and other respiratory infections *. Rev Lat Am Enfermagem. 28:e3353.

6. Hao W, Xu G, Wang Y. Factors influencing the filtration performance of homemade face masks. Journal of Occupational and Environmental Hygiene. 2021 Mar 4;18(3):128–38.

7. Konda A, Prakash A, Moss GA, Schmoldt M, Grant GD, Guha S. Aerosol Filtration Efficiency of Common Fabrics Used in Respiratory Cloth Masks. ACS Nano. 2020 Apr 24;acsnano.0c03252.

8. Smith JD, MacDougall CC, Johnstone J, Copes RA, Schwartz B, Garber GE. Effectiveness of N95 respirators versus surgical masks in protecting health care workers from acute respiratory infection: a systematic review and meta-analysis. CMAJ. 2016 May 17;188(8):567–74.

9. Long Y, Hu T, Liu L, Chen R, Guo Q, Yang L, et al. Effectiveness of N95 respirators versus surgical masks against Influenza: A systematic review and meta-analysis. J Evid Based Med. 2020 May;13(2):93–101.

10. Sakai N, Yoshido J, Kinjo S. Improved Polyethylene Terephthalate shield and the mitigation of COVID-19 aerosols during airway manipulation. J Case Rep. 2021;1(1):1001.

11. van Doremalen N, Bushmaker T, Morris DH, Holbrook MG, Gamble A, Williamson BN, et al. Aerosol and Surface Stability of SARS-CoV-2 as Compared with SARS-CoV-1. New England Journal of Medicine. 2020 Apr 16;382(16):1564–7.

12. Guo L, Wang M, Zhang L, Mao N, An C, Xu L, et al. Transmission risk of viruses in large mucosalivary droplets on the surface of objects: A time-based analysis. Infect Dis Now. 2021 May;51(3):219–27.

13. Hewawaduge C, Senevirathne A, Jawalagatti V, Kim JW, Lee JH. Copper-impregnated three-layer mask efficiently inactivates SARS-CoV2. Environ Res. 2021 May;196:110947.

14. Li Y, Leung P, Yao L, Song QW, Newton E. Antimicrobial effect of surgical masks coated with nanoparticles. Journal of Hospital Infection. 2006;62(1):58–63.

15. Rai M, Deshmukh SD, Ingle AP, Gupta IR, Galdiero M, Galdiero S. Metal nanoparticles: The protective nanoshield against virus infection. Crit Rev Microbiol. 2016;42(1):46–56.

16. Maduray K, Parboosing R. Metal Nanoparticles: a Promising Treatment for Viral and Arboviral Infections. Biol Trace Elem Res. 2021 Aug;199(8):3159–76.

17. Shirvanimoghaddam K, Akbari MK, Yadav R, Al-Tamimi AK, Naebe M. Fight against COVID-19: The case of anti-viral surfaces. APL Materials. 2021 Mar;9(3):031112.

18. Angelé-Martínez C, Nguyen KVT, Ameer FS, Anker JN, Brumaghim JL. Reactive oxygen species generation by copper(II) oxide nanoparticles determined by DNA damage assays and EPR spectroscopy. Nanotoxicology. 2017 Mar;11(2):278–88.

19. Jeremiah SS, Miyakawa K, Morita T, Yamaoka Y, Ryo A. Potent anti-viral effect of silver nanoparticles on SARS-CoV-2. Biochem Biophys Res Commun. 2020 Nov 26;533(1):195–200.

20. Reinhold D, Brocke S. The differential roles of zinc in immune responses and their potential implications in anti-viral immunity against SARS-CoV-2. Clin Nutr. 2021 Feb;40(2):652.

21. Velthuis AJW te, Worm SHE van den, Sims AC, Baric RS, Snijder EJ, Hemert MJ van. Zn2+ Inhibits Coronavirus and Arterivirus RNA Polymerase Activity In Vitro and Zinc Ionophores Block the Replication of These Viruses in Cell Culture. PLOS Pathogens. 2010 Nov 4;6(11):e1001176.

22. Xiang D xi, Chen Q, Pang L, Zheng C long. Inhibitory effects of silver nanoparticles on H1N1 Influenza A virus in vitro. J Virol Methods. 2011 Dec;178(1–2):137–42.

23. Hodek J, Zajícová V, Lovëtinská-Šlamborová I, Stibor I, Müllerová J, Weber J. Protective hybrid coating containing silver, copper and zinc cations effective against human immunodeficiency virus and other enveloped viruses. BMC Microbiology. 2016 Apr 1;16(1):56.

24. Manuel CS, Moore MD, Jaykus LA. Efficacy of a disinfectant containing silver dihydrogen citrate against GI.6 and GII.4 human norovirus. J Appl Microbiol. 2017 Jan;122(1):78–86.

25. Chen YN, Hsueh YH, Hsieh CT, Tzou DY, Chang PL. Anti-viral Activity of Graphene–Silver Nanocomposites against Non-Enveloped and Enveloped Viruses. Int J Environ Res Public Health. 2016 Apr;13(4):430.

26. Firquet S, Beaujard S, Lobert PE, Sané F, Caloone D, Izard D, et al. Survival of Enveloped and Non-Enveloped Viruses on Inanimate Surfaces. Microbes Environ. 2015 Jun;30(2):140–4.

27. Vasickova P, Pavlik I, Verani M, Carducci A. Issues Concerning Survival of Viruses on Surfaces. Food Environ Virol. 2010;2(1):24–34.

28. Chandran K, Farsetta DL, Nibert ML. Strategy for non-enveloped virus entry: a hydrophobic conformer of the reovirus membrane penetration protein micro 1 mediates membrane disruption. J Virol. 2002 Oct;76(19):9920–33.

29. Kaufman SS, Green KY, Korba BE. Treatment of norovirus infections: moving anti-virals from the bench to the bedside. Antiviral Res. 2014 May;105:80–91.

30. Tremiliosi GC, Simoes LGP, Minozzi DT, Santos RI, Vilela DCB, Durigon EL, et al. Ag nanoparticles-based anti-microbial poly-cotton fabrics to prevent the transmission and spread of SARS-CoV-2 [Internet]. bioRxiv; 2020 [cited 2022 May 26]. Available from: https://www.biorxiv.org/content/10.1101/2020.06.26.152520v1

31. Borkow G, Zhou SS, Page T, Gabbay J. A Novel Anti-Influenza Copper Oxide Containing Respiratory Face Mask. PLOS ONE. 2010 Jun 25;5(6):e11295.

32. Hamouda T, Ibrahim HM, Kafafy HH, Mashaly HM, Mohamed NH, Aly NM. Preparation of cellulose-based wipes treated with anti-microbial and anti-viral silver nanoparticles as novel effective high-performance coronavirus fighter. International Journal of Biological Macromolecules. 2021 Jun 30;181:990–1002.

33. Qiao Y, Yang M, Marabella IA, McGee DAJ, Aboubakr H, Goyal S, et al. Greater than 3-Log Reduction in Viable Coronavirus Aerosol Concentration in Ducted Ultraviolet-C (UV–C) Systems. Environ Sci Technol. 2021 Apr 6;55(7):4174–82.

34. Huber T, Goldman O, Epstein AE, Stella G, Sakmar TP. Principles and practice for SARS-CoV-2 decontamination of N95 masks with UV-C. Biophys J. 2021 Jul 20;120(14):2927–42.

35. Hasani M, Campbell T, Wu F, Warriner K. Decontamination of N95 and surgical masks using a treatment based on a continuous gas phase-Advanced Oxidation Process. PLOS ONE. 2021 Mar 18;16(3):e0248487.

36. Liu DX, Liang JQ, Fung TS. Human Coronavirus-229E, -OC43, -NL63, and -HKU1 (Coronaviridae). Encyclopedia of Virology. 2021;428–40.

37. Vijayan P P, P.g C, Abraham P, George JS, Maria HJ, T S, et al. Nanocoatings: Universal antiviral surface solution against COVID-19. Progress in Organic Coatings. 2022 Feb 1;163:106670.

38. Balagna C, Perero S, Percivalle E, Nepita EV, Ferraris M. Virucidal effect against coronavirus SARS-CoV-2 of a silver nanocluster/silica composite sputtered coating. Open Ceramics. 2020 May;1:100006.

39. Freymuth F, Vabret A, Rozenberg F, Dina J, Petitjean J, Gouarin S, et al. replication of respiratory viruses, particularly influenza virus, rhinovirus, and coronavirus in HuH7 hepatocarcinoma cell line. Journal of Medical Virology. 2005;77(2):295–301.

40. Mathew E, Gilmore BF, Larrañeta E, Lamprou DA. Antimicrobial 3D Printed Objects in the Fight Against Pandemics. 3D Printing and Additive Manufacturing. 2021 Feb;8(1):79–86.

41. Pollard ZA, Karod M, Goldfarb JL. Metal leaching from anti-microbial cloth face masks intended to slow the spread of COVID-19. Sci Rep. 2021 Sep 28;11(1):19216.

42. Ng CT, Yong LQ, Hande MP, Ong CN, Yu LE, Bay BH, et al. Zinc oxide nanoparticles exhibit cytotoxicity and genotoxicity through oxidative stress responses in human lung fibroblasts and Drosophila melanogaster. Int J Nanomedicine. 2017 Feb 28;12:1621–37.

